# Modeling a gene regulatory network of EMT hybrid states for mouse embryonic skin cells

**DOI:** 10.1101/799908

**Authors:** Dan Ramirez, Vivek Kohar, Ataur Katebi, Mingyang Lu

**Affiliations:** College of Health Solutions, Arizona State University, Tempe, Arizona, United States of America; The Jackson Laboratory, Bar Harbor, Maine, United States of America

## Abstract

Epithelial-mesenchymal transition (EMT) plays a crucial role in embryonic development and tumorigenesis. Although EMT has been extensively studied with both computational and experimental methods, the gene regulatory mechanisms governing the transition are not yet well understood. Recent investigations have begun to better characterize the complex phenotypic plasticity underlying EMT using a computational systems biology approach. Here, we analyzed recently published single-cell RNA sequencing data from E9.5 to E11.5 mouse embryonic skin cells and identified the gene expression patterns of both epithelial and mesenchymal phenotypes, as well as a clear hybrid state. By integrating the scRNA-seq data and gene regulatory interactions from the literature, we constructed a gene regulatory network model governing the decision-making of EMT in the context of the developing mouse embryo. We simulated the network using a recently developed mathematical modeling method, named RACIPE, and observed three distinct phenotypic states whose gene expression patterns can be associated with the epithelial, hybrid, and mesenchymal states in the scRNA-seq data. Additionally, the model is in agreement with published results on the composition of EMT phenotypes and regulatory networks. We identified Wnt signaling as a major pathway in inducing the EMT and its role in driving cellular state transitions during embryonic development. Our findings demonstrate a new method of identifying and incorporating tissue-specific regulatory interactions into gene regulatory network modeling.

**Author Summary:** Epithelial-mesenchymal transition (EMT) is a cellular process wherein cells become disconnected from their surroundings and acquire the ability to migrate through the body. EMT has been observed in biological contexts including development, wound healing, and cancer, yet the regulatory mechanisms underlying it are not well understood. Of particular interest is a purported hybrid state, in which cells can retain some adhesion to their surroundings but also show mesenchymal traits. Here, we examine the prevalence and composition of the hybrid state in the context of the embryonic mouse, integrating gene regulatory interactions from published experimental results as well as from the specific single cell RNA sequencing dataset of interest. Using mathematical modeling, we simulated a regulatory network based on these sources and aligned the simulated phenotypes with those in the data. We identified a hybrid EMT phenotype and revealed the inducing effect of Wnt signaling on EMT in this context. Our regulatory network construction process can be applied beyond EMT to illuminate the behavior of any biological phenomenon occurring in a specific context, allowing better identification of therapeutic targets and further research directions.

## Introduction

Epithelial-mesenchymal transition is a widely studied cellular process during which epithelial cells lose the junctions binding them to their immediate environment while simultaneously acquiring the phenotypic traits of mesenchymal cells, which permit migratory and invasive behaviors [1,2]. There are three distinct types of EMT in the contexts of embryonic development, wound healing, and cancer progression [3]. One major topic of interest regarding EMT is the stability, structure, and function of hybrid phenotypes [4], in which cells express canonical markers of both epithelial (E) and mesenchymal (M) phenotypes. However, it is still unclear whether such hybrid phenotypes are merely transitional states or a distinct hybrid cell type [5,6]. A hybrid phenotype in cancer could permit the formation of circulating tumor cell clusters, groups of cells which can collectively migrate, increasing their likelihood of successfully forming a secondary tumor [7]. On the other hand, partial EMT phenotypes may be helpful in their ability to collectively migrate and close open wounds [3,8]. A greater understanding of the mechanisms of EMT with respect to these hybrid cells could therefore permit more advanced investigation and treatment options in a number of clinical situations.

To better understand the regulatory mechanisms that control the creation and maintenance of these cell types and the dynamical transitions between them, researchers have adopted systems-biology approaches to model the gene regulatory networks (GRNs) that govern the decision-making of EMT [9–15]. A number of simple gene regulatory circuit models have been proposed which would permit the existence of three or more states during EMT based on the activity of core transcription factors (TFs) including Snail and Zeb, as well as other regulatory elements such as microRNAs [14–17]. Beyond these reduced models, larger networks have been simulated to observe the abilities of different signal transduction pathways to induce and regulate EMT [9,10]. Building on the large body of experimental evidence for specific gene regulatory interactions, GRN models can be constructed which accurately convey the general phenotypic topography of EMT [17,18]. However, such methods are usually limited by insufficient experimental evidences on regulatory interactions and human errors in the process of curation. Moreover, literature-based GRNs are often composed of interactions identified in different contexts; therefore, it is difficult to draw biologically relevant conclusions for specific systems.

While many of the above-mentioned approaches use experimental data on specific biomarkers to validate their models, with the advent of new genomics technologies, it is now possible to measure genome-wide transcriptomics data for different stages of the process. Especially with single cell measurement, one can investigate the heterogeneity of a cell population and distinguish between stable hybrid phenotypes and simple mixtures of E and M cells. A 2018 publication by Dong et al. performed an analysis of EMT in 1916 embryonic mouse cells, demonstrating the presence of three distinct phenotypes in the data and examining the relationship between EMT, stemness, and developmental pseudotime [5]. Using the complete gene expression profiles of each individual cell may allow: (1) far more thorough and conclusive investigations into the presence of a hybrid phenotype; (2) discovery of important biological signaling pathways that drive EMT; (3) inference of a GRN model directly from transcriptomics data via computational algorithms, such as SCENIC and metaVIPER [13,19–21]. These algorithms consider metrics such as coexpression patterns, and TF binding motifs to build complete GRNs from experimental data. Unfortunately, it remains challenging to build GRNs directly from experimental gene expression data (1) to recapitulate causal regulatory links and (2) to elucidate the dynamical behavior of a biological system.

Here, we aimed to bridge the gap between the top-down genomics approaches and the bottom-up literature-based approaches to understand the context specific gene regulatory mechanisms of EMT. We explore the option to start from a literature-based network, and then refine it to reflect the TF-target relationships specific to a scRNA-seq dataset on E9.5 to E11.5 mouse embryonic skin cells. We combined a SCENIC analysis of the expression data with published information regarding the EMT regulatory network to develop a small gene network model which predicts several distinct states during EMT similar to those observed in the scRNA-seq data. We then annotated the phenotypes as epithelial, mesenchymal, and hybrid by comparing gene expression profiles with canonical markers and well-documented evidence regarding the composition of each phenotype. From the scRNA-seq data, we identified Wnt as the most active signaling pathway regulating EMT in this context and modeled its effect on the distribution of phenotypic states. This application of modeling techniques in combination with single-cell data allows the construction of highly representative models and accurate predictions regarding the phenotype of cells in a specific context.

## Results

### scRNA-seq data identifying hybrid EMT states

We first analyzed public scRNA-seq data from three mouse embryos at three different developmental stages ranging from E9.5-E11.5 [5]. Of the eight tissue types sequenced in the dataset, we examined the four for which cells were designated as epithelial (E) or mesenchymal (M) according to the location in the embryo from which they were collected, namely lung, liver, skin, and intestine. Because the cells separated according to phenotype most clearly in skin cells and because the skin cells provided the most evidence for co-expression of E and M marker genes (Fig. S1), suggesting a prevalent hybrid state, we chose to focus specifically on the 156 skin cells for further analysis.

To evaluate the overall structure of the data, PCA was performed on the log-normalized unique molecular identifier (UMI) counts for all 16082 genes in the skin cell dataset. This dimensional reduction showed the cells to be immediately distinguishable according to their phenotype, with epithelial (E) and mesenchymal (M) cells forming independent groups which could be identified by density-based clustering (Fig. 1a, cell type annotated by color). The first two principal components effectively separate E from M cells, indicating robust phenotypic differences between the cell types. Cells of the same developmental stages tended to appear alongside each other (Fig. 1a, stages denoted by point shape), suggesting a noticeable developmental bias in the dimensional reduction.

**Figure 1.**
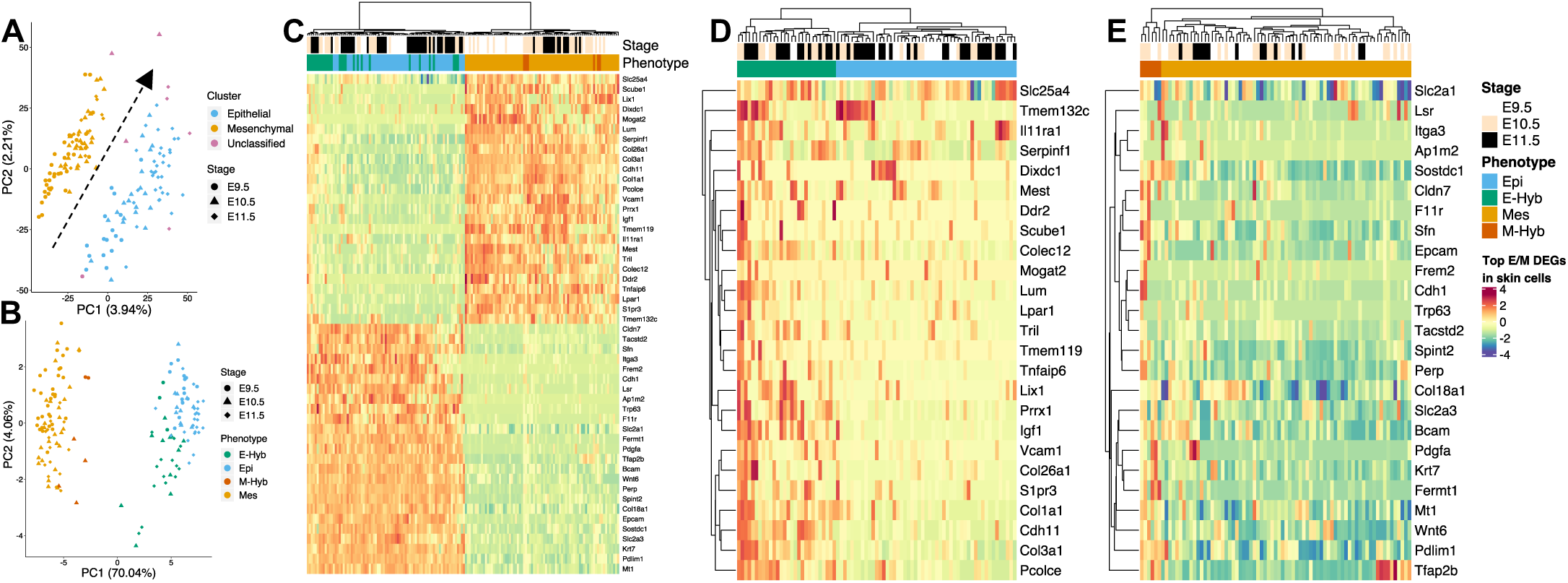
Transcriptional analysis of 156 embryonic mouse skin cells, stages E9.5-E11.5. (A) PCA on all 16082 genes with density-based clustering. PC1 captures ∼ 4% of the variance and effectively separates E and M phenotypes. Note clear developmental trends on illustrated diagonal. (B) PCA on top 25 E and 25 M marker genes as identified by DEG analysis. PC1 captures ∼ 70% of the variance and clearly separates E and M clusters, while hybrid cell populations appear nearer to the center. (C) Heatmap on top 50 EMT-related genes color coded by cluster. Hierarchical clustering groups E and M cells together, with E-Hyb and M-Hyb forming distinct subclusters with discernible co-expression. (D) Heatmap of E cells using top 25 M marker genes. Hierarchical clustering separates the population into two distinct subclusters marked by high and low co-expression of M markers, denoted E-Hyb and E. (E) Heatmap of M cells using top 25 E markers. Once again hierarchical clustering separates two subpopulations with higher and lower levels of co-expression.

To focus specifically on the changes associated with EMT independent of development, differential expression analysis was performed on the E and M clusters, providing a set of epithelial and mesenchymal markers relevant to this biological context. Among the most differentially expressed genes (DEGs) were known EMT markers including E-cadherin (Cdh1), epithelial cellular adhesion molecule (Epcam), keratin 7 (Krt7), and collagen type I alpha 1 chain (Col1a1) [22–24]. We then performed PCA on the top 50 EMT-related DEGs (Fig. 1b) to examine the distribution of cells on the EMT phenotypic landscape. The first principal component here captured nearly 68% of the variance in the data, clearly separating the E and M clusters as denoted by the previous study. The PCA also showed dramatically less developmental bias in these results, suggesting independent mechanisms of development and EMT at work in the cells.

The abovementioned PCA indicated the presence of several distinct EMT states in the data, but the sharp contrast between E and M cells obscured subtler differences which could mark a hybrid state. Hierarchical clustering was performed on the expression values of the top 25 marker DEGs for each cluster (Fig. 1c). While the expression heatmap shows distinct regions of coexpression, once again the less dramatic gene expression profile of the hybrid cells was overshadowed by the greater contrast between E and M cells. Because a previous investigation [5] of the same dataset uncovered hybrid cells only as a subpopulation of the E cells, the data were split into separate groups of E and M cells and PCA was performed on the top 25 marker DEGs of the other cluster; i.e. E cells were clustered on the top 25 M genes and M cells on the top 25 E genes (Fig. 1d-e). Among the E cells, a clear subpopulation was discernible with higher expression of M marker genes, suggesting a hybrid phenotype. These cells, designated by the dendrogram on **Fig. 1d** (cut at the number of clusters indicated by the Ball Index [25]), were denoted E-Hybrid (E-Hyb). The same approach for the M cells yielded fewer M-Hybrid (M-Hyb) cells, but this smaller subpopulation also showed unusually high expression of E marker genes, indicating that a M-Hyb phenotype may be present in small quantities. Overall, the DEG analysis identified multiple EMT states in the data including hybrid states, suggesting that EMT is occurring in embryonic mouse tissues between the E9.5-E11.5 stages.

We also examined the distribution of the EMT states across developmental stages using the scRNA-seq. However, the data show few well-distinguished trends in the relative proportions of different phenotypes across the E9.5-E11.5 timepoints, counter to the gradual increase in M cells that might be expected if EMT were occurring (Fig. S2). Previous experiments have found that EMT does occur in the developmental mouse during these stages but have not established with certainty the direction or the volume of EMT which occurs [2,26,27]. It is also hard to identify such information with the scRNA-seq data, likely because of low sample size of single cells and the lack of time-series data.

### Constructing a gene regulatory network for EMT

To create a GRN which is both relevant to the specific dataset in this study and representative of the regulatory mechanisms of EMT in general, we devised a computational protocol to incorporate interactions from both literature and the scRNA-seq data analysis (Fig. 2a). Beginning from a literature-based network (Fig. 2a, leftmost diagram), we removed genes that are vastly not expressed and signaling pathways (second diagram) to identify a small set of core regulators. Then, using gene-set enrichment analysis (GSEA) on experimental data, we reincorporated the most enriched signaling pathway as an upstream driver of the network (third diagram, yellow node and edges). Finally, using SCENIC, we inferred the regulatory activity of the TFs in the dataset and introduced context-specific interactions (rightmost diagram, green nodes and edges) to generate the network to be simulated using mathematical modeling (see below for details).

**Figure 2.**
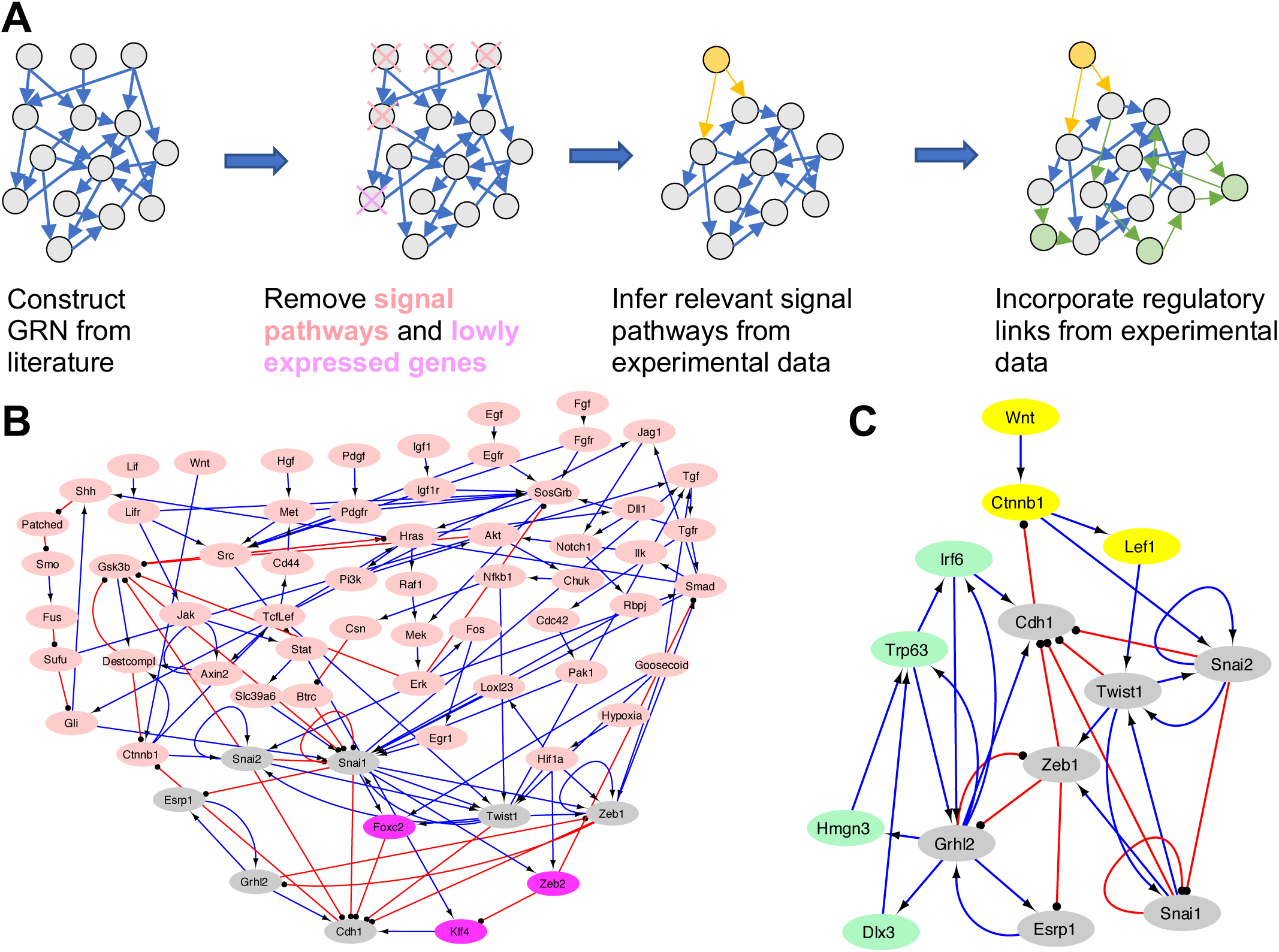
Construction of an EMT gene regulatory network that integrates scRNA-seq data. (A) Flowchart depicting GRN construction process. A GRN was built based on information on EMT in the literature and subsequently filtered to remove lowly expressed genes and signaling pathways. The signaling pathway(s) of interest are then implemented based on information from GSEA and additional regulatory links and nodes are incorporated based on the SCENIC results. (B) Literature-based network with nodes removed color coded in red and purple. (C) GRN after incorporating links from SCENIC, with 14 nodes and 34 edges.

We first started from a 66-node and 130-edge gene regulatory network from previous experimental and GRN modeling studies on EMT (Fig. 2b, Tables S1-2). To focus specifically on the core gene regulatory interactions that are present in the current data set, the network was adjusted to remove signaling pathways and genes with consistently low expression. Because Wnt signaling was indicated to be most relevant in the pathway enrichment analysis (details below), it was reintroduced to the network as a direct activating signal towards b-catenin (Ctnnb1), which in turn activates Lef1 and Snai2 and is inhibited by Cdh1 [10,28]. From the above-mentioned procedures, we constructed an initial 10-node GRN, as shown in Fig. 2c, gray and yellow nodes. A central feature of the network is a triangular interaction between Grhl2, Zeb1, and Cdh1, in which Zeb1 and Grhl2 exhibit mutual inhibition and Zeb1 inhibits Cdh1, while Grhl2 activates Cdh1. Both Zeb and Grhl have been extensively studied as important regulators of EMT [14,29,30].

To further improve the initial GRN to reflect additional new context-specific interactions in the dataset of interest, we applied SCENIC to infer additional regulatory genes and links from the scRNA-seq data. Here, using the 10 genes in the initial core network, we collected from SCENIC any new regulatory link in which either the regulator or the targeted gene is already in the network (all first neighbor nodes). These first-neighbor interactions were further filtered based on mean regulon activity across cell types, such that the only interactions kept were those within the top 25 most differentially active regulons for E and M cells. Autoregulating interactions were also removed, as well as genes with consistently low regulatory activity as inferred by SCENIC. We further removed genes which were not TFs from the network, resulting in a model of 26 nodes and 79 edges. Finally, we removed interactions from SCENIC which were not supported by expression or activity data, resulting in the final network of 14 nodes and 34 edges (see methods) (Fig. 2c).

Of the four genes added to the network, namely interferon regulatory factor 6 (Irf6), transformation related protein 63 (Trp63), high mobility group nucleosome-binding protein 3 (Hmgn3), and distal-less homeobox 3 (Dlx3), all were closely integrated with the E TFs in the network, primarily Grhl2. Both upstream and downstream interactions were incorporated, with many serving to directly or indirectly activate Cdh1. Some of the genes added to the network during the process have been studied in relation to EMT before, such as *Trp63* [31], and *Irf6* [32]. The contribution of the genes in the GRN to EMT in embryonic mouse tissues is also supported by previous gene expression experiments (Table S3).

### Identifying the role of Wnt signaling in network dynamics

To better understand the signaling pathways involved in regulating EMT in this context, the full list of DEGs between E and M cells from Seurat were supplied as input to enrichR, an R package which performs enrichment analysis on a list of genes. EnrichR examined the list of genes to determine which of 303 KEGG 2019 pathways for *Mus musculus* were overrepresented. The Hippo signaling pathway was the most prevalent signaling pathway among the results, overrepresented in E cells with an adjusted p-value <0.05 and combined score of 37.7 (Table 1). Among the leading-edge genes in the top enriched pathways are several genes in the Wnt signaling gene family. Moreover, there is substantial crosstalk between the Wnt/b -catenin and Hippo pathways [33]. A second gsea analysis was conducted using fgsea, an R package which uses a ranked list of genes, with the average log fold change across clusters as the ranking metric. This analysis also found the Wnt and Hippo signaling pathways to be enriched in E cells, albeit with lower levels of significance (Table S4). Because Wnt signaling was identified as discernibly enriched in the EMT process, we chose to further investigate the role of Wnt in driving EMT by network modeling.

**Table 1:** Top 10 EMT related signaling pathways identified by enrichR sorted by combined score.

| Pathway | Adjusted p-value | Odds Ratio | Combined Score |
| --- | --- | --- | --- |
| Hippo signaling pathway | 0.0003 | 3.2037 | 37.6884 |
| Wnt signaling pathway | 0.0008 | 3.0161 | 31.3515 |
| PI3K-Akt signaling pathway | 0.0004 | 2.328 | 26.2993 |
| Relaxin signaling pathway | 0.0053 | 2.8652 | 22.5057 |
| AGE-RAGE signaling pathway in diabetic complications | 0.0133 | 2.9199 | 19.274 |
| Rap1 signaling pathway | 0.0098 | 2.309 | 16.2581 |
| p53 signaling pathway | 0.0366 | 3.0208 | 16.1064 |
| Estrogen signaling pathway | 0.0351 | 2.4009 | 13.0239 |
| Ras signaling pathway | 0.0469 | 1.9561 | 9.8028 |
| mTOR signaling pathway | 0.0846 | 2.0891 | 9.0981 |

The expression patterns of genes in the Wnt signaling pathway were also examined with pathview [34], which generates color coded diagrams to reflect the activity of KEGG pathways in a dataset of interest. As shown in **Fig. 3**, Wnt signaling is more active in the E cell population than that in the M cell population. Genes are up- and down-regulated in accordance with the currently known regulatory interactions in the Wnt pathway, suggesting that Wnt signaling is substantially active in the E cells in this dataset. Together, the simulation and expression data are a compelling indication of the role of Wnt signaling in inducing EMT.

**Figure 3.**
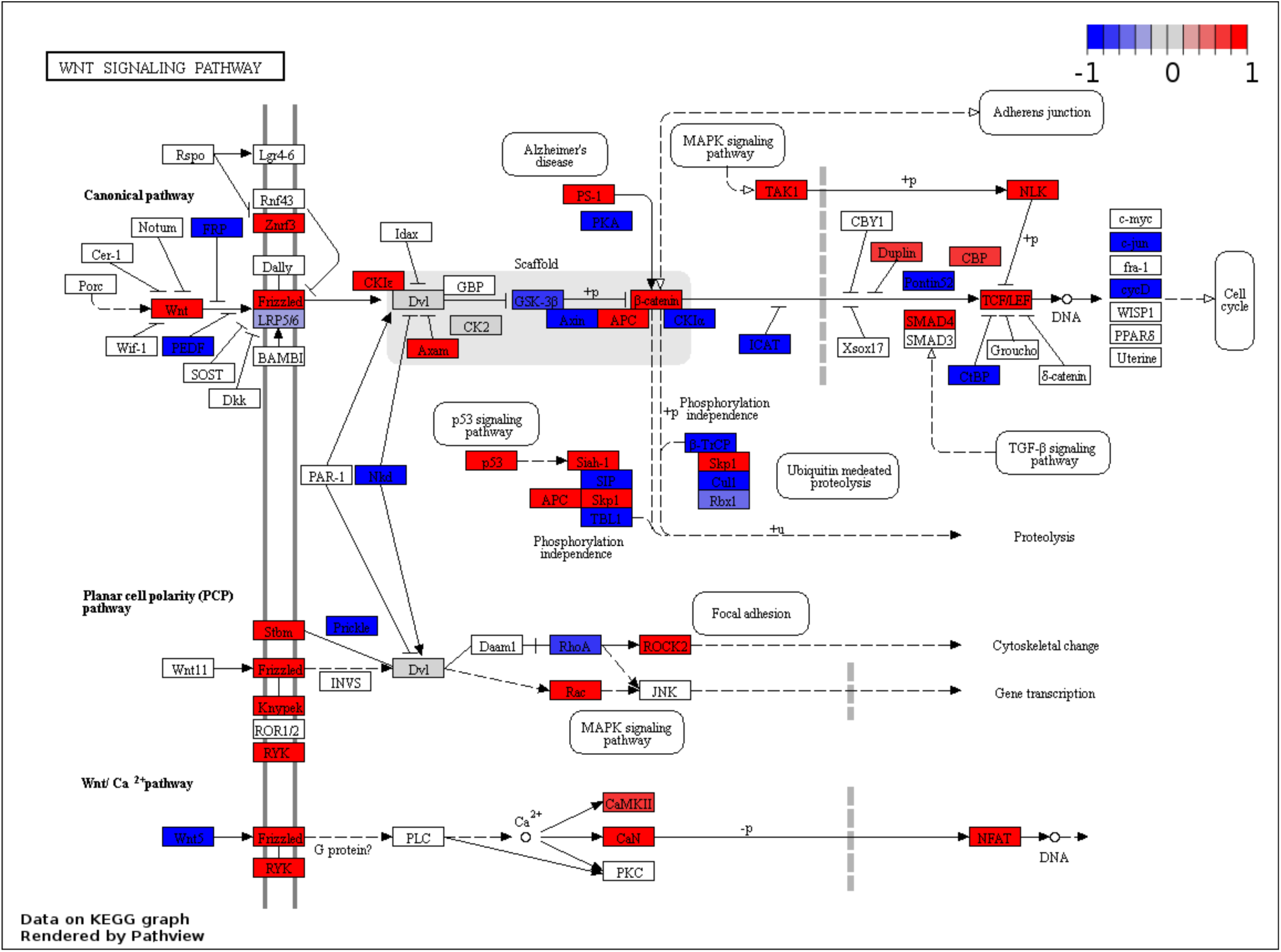
The role of Wnt signaling in regulating EMT. Color-coded KEGG pathway for Wnt signaling in Mus musculus based on t-statistics from GSEA analysis. Color coding scheme represents the expression profile of E cells, with red signifying high expression and blue signifying low expression. Genes not present in the dataset have no background fill.

### Network dynamics are consistent with the scRNA-seq data

To evaluate the dynamical behavior of the 14-node GRNs, we applied RACIPE, a mathematical modeling algorithm, (see methods for details) to generate simulated gene expression profiles from an ensemble of 10,000 models with randomly generated parameters. Using stochastic analysis and simulated annealing, we modeled the network at 30 progressively smaller noise levels, capturing the relative stability of states through their prevalence in the simulation results. We applied hierarchical clustering to identify three groups in the simulated gene expression profiles, which could easily be identified as epithelial, hybrid, and mesenchymal phenotypes (Fig. 4a). The latter three phenotypes agreed in general with the established understanding of EMT phenotypes. Models corresponding to the E phenotype, accounting for approximately 40.8% of the models, showed high expression of Cdh1, Grhl2, Esrp1, and several other TFs involved in positive feedback loops with the E marker genes. Models describing the M state, comprising 37.7% of the total, showed high expression of Zeb1, Twist1, Snai2, and other M markers. These expression profiles are largely consistent with previous research on EMT and many of the genes which identify phenotypes in the simulations are commonly used marker genes in experiments, such as Cdh1, Zeb1, and Twist1 [9,10,35,36]. The hybrid models, which comprised the remaining 21.5% of the total expressed all genes in the network to some extent, though Cdh1 and Zeb1 had lower levels. Overall, the simulation agreed with our prediction that the network permits two distinct states and a third hybrid state defined by coexpression of E and M markers.

**Figure 4.**
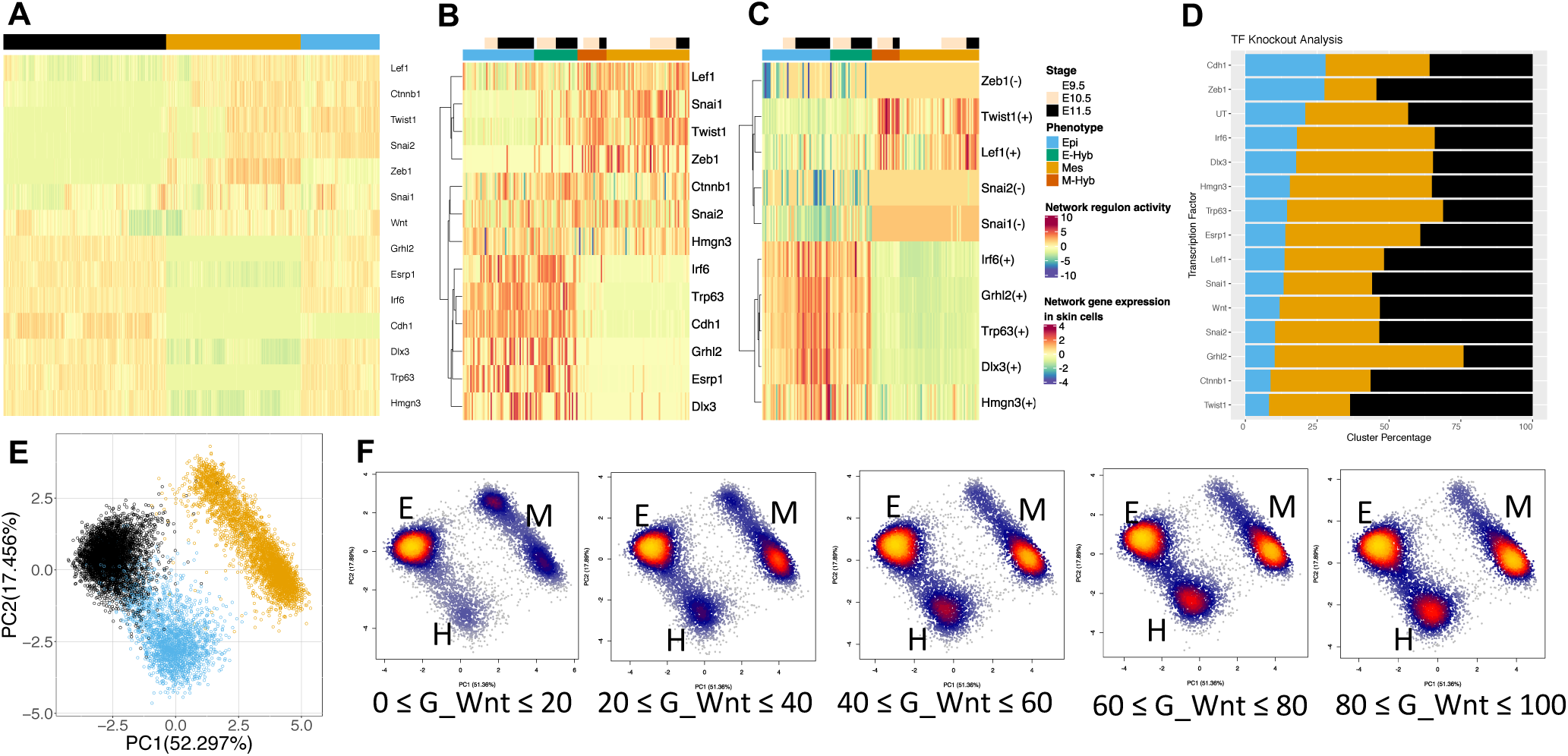
Mathematical modeling of the constructed EMT gene network. (A) Heatmap of RACIPE simulation results for 14-node GRN with 3 most prevalent phenotypes identified via hierarchical clustering. E, M and H phenotypes are discernible by expression of noted marker genes and co-expression. (B) Heatmap of inferred activity of regulons present in the GRN in skin cells. Note simultaneous activity of E and M regulons in hybrid cell types (C) Heatmap of network gene expression in skin cells. E and M cells show high expression of their respective marker genes and hybrid cells show coexpression of both. (D) Knockdown subset analysis of RACIPE results sorted by resulting prevalence of H models. Untreated condition (UT) represents the normal simulations without knocking down. (E) PCA of simulated network gene expression values color coded by cluster. (F) Results from Wnt perturbation simulations projected onto the first two principal component axes of the original simulation. Wnt production rate increases from left to right in increments of 20% of the maximum parameter value. Clusters correspond to the labeled phenotypic states. As Wnt signaling increases, the E state generally decreases in prevalence, while the H and M states increase.

In addition to the stochastic simulations, we conducted deterministic simulations of the 14-node network. The deterministic simulations generated the same three phenotypes as well as a number of models with low expression for all genes (a low-expression state, Fig. S3). With respect to the distribution of phenotypes, the deterministic simulations yielded a greater proportion of M models and fewer E and H models, but overall the results were comparable. The results are also consistent with our previous studies that stochastic analysis yields less of the low-expression state than the deterministic analysis [9,37] Because the stochastic analysis allows better evaluation of the stability of various states better, we proceeded with the stochastic simulation results.

We also compared the simulation results with stochastic RACIPE simulations of the core network prior to incorporating interactions from SCENIC (Fig. S4). The three phenotypes present aligned closely with the phenotypes predicted with the core network, although the updated network topology resulted in a larger proportion of H models. This suggests that the fundamental behaviors of EMT are highly conserved across biological contexts and tissue-specific interactions account for small optimizations. One of the potential roles of these genes newly added to the network is to stabilize a hybrid phenotype during development but they may not be involved in other contexts such as wound healing.

To validate the GRN model, we compared the simulated gene expression profiles with scRNA-seq data (Fig. 4b). The gene expression data aligned well with the simulation results, with E cells showing high expression of the E markers predicted by RACIPE, M cells showing expression of M markers, and E-Hyb and M-Hyb cells showing coexpression of E and M markers. This alignment suggests that the network accurately captured the behavior of EMT in the context of this dataset. The M-Hyb cells showed weaker coexpression, likely because of the small number of cells. Generally, the single cell expression data can be grouped into the same three main clusters present in the simulation data. The genes involved in Wnt signaling, namely Ctnnb1 and Lef1, are less effective markers for the different phenotypes, likely due to the complex mechanics of signal transduction and the influences not captured by gene expression alone.

To consider the cases where TF expression does not correlate with TF activity, we inferred the regulon activity for each TF using the expression of targeted genes (Fig. 4c). Epithelial and E-Hyb cells show strong agreement with the RACIPE results, with high activity in Irf6, Grhl2, Trp63, Dlx3, and Hmgn3. Similarly, the M cells show high activity among M marker TFs including Twist1, Zeb1, and Snai1. The hybrid cells showed increased activity in Snai2 and Lef1 in particular, with some activity in the other M marker TFs, generally in agreement with the simulations. Snai2 and Hmgn3 show activity profiles dramatically different from their expression profiles, probably because expression is not always indicative of TF activity. The three states found in the RACIPE simulations are also observable using regulon activity, although using a larger number of TFs would likely facilitate the identification of the hybrid state.

We examined the distribution of phenotypes in two-dimensional space using PCA of the simulated gene expression values (Fig. 4e). This analysis revealed that the H models grouped more closely to the E models than the M models, indicating that the hybrid state may be more closely related to the E phenotype. This may also reflect the fact that more E-Hyb cells were identified in the scRNA-seq data because SCENIC may have identified primarily interactions supporting an E-Hyb phenotype. The M models also formed a less centralized cluster on the PCA plot, suggesting there may be more phenotypic variety among M cells than E cells with respect to genes involved in EMT.

The effects of perturbations on specific genes on the network were examined through subsequent knockdown simulations. The proportions of each phenotype with a gene knockdown were compared to the proportions for the untreated conditions (Fig. 4d). Grhl2 and Zeb1 had notable effects when knocked down, reducing the proportion of E and M cells respectively, accurately reflecting their central positions in the network topology as well as the mutual inhibition between them. Knockdown of Wnt also appears to influence the phenotypic distribution, resulting in fewer H and M cells and more E cells. The same effect is present to a greater degree when knocking down the direct downstream target of Wnt, Ctnnb1, suggesting that Wnt signaling plays a role in driving EMT and potentially in inducing the H phenotype. Our results on the perturbation of Wnt signaling are consistent with experimental findings that Wnt signaling can induce EMT and may be implicated in cancer metastasis as well [28,38,39].

The genes incorporated into the network from SCENIC all showed similar impacts when knocked down, reducing the proportion of E cells and increasing the prevalence of M cells. These knockdowns also precipitated a decrease in the number of H cells, suggesting that the hybrid phenotype is regulated by a combination of E and M TFs. Zeb1 and Cdh1 are unique in that knockdowns to these genes increase the prevalence of the H state, likely reflecting the negative feedback which heavily influences both of these genes in the topology. They are also the only two genes which are not strongly expressed in the H state, indicating they strongly influence the network in favor of the M and E states, respectively. Because the genes are so central to the regulatory landscape of the M and E phenotypes respectively, when their production is knocked down, the phenotypic distribution shifts in favor of states which are characterized by intermediate or low expression of them.

Wnt expression alone is a notably poor determinant of phenotype in the RACIPE results because of its integration into the network topology; as an input to the system, it has no regulating influences other than the randomly generated kinetic parameters and thus would be expected to show unpredictable gene expression values. However, as shown by the direct downstream target of Wnt, Ctnnb1, this effect attenuates almost immediately and the influence of Wnt signaling can be seen through the genes with which it indirectly interacts.

To further examine the role of Wnt signaling in the EMT process, perturbation simulations were performed with five different ranges of Wnt production rates from low to high (Fig. 4f). As the level of Wnt expression increases, the E cluster generally diminishes while the M and H clusters grow. This indicates that Wnt signaling in the network serves to promote M and H phenotypes by inducing EMT.

## Discussions

Mathematical modeling of GRNs has traditionally been conducted using information from published literature, which is limited by experimental data that is too noisy or ambiguous to neatly reflect the predictions of the model. Additionally, there are many difficulties of integrating previous results from disparate sources. The approach developed here addresses these limitations by building upon a regulatory network which is well supported in a number of biological contexts and elucidating the specific interactions at work in a particular dataset. Using DEG analysis and GSEA, we were able to identify different phenotypes in scRNA-seq data and illuminate the activity of different signaling pathways. Using SCENIC, we characterized the regulatory networks present in the data and incorporated this information into a literature-based network modeling EMT. The states predicted by our simulations are in agreement with both previous results and the single-cell expression data, suggesting the mechanics of EMT in this context are well represented in the network topology. We are able to clearly identify three distinct expression patterns using the genes in the network, correlating well with general understanding of E, M, and E/M hybrid cells. Furthermore, the perturbation simulations provide potential directions for the development of interventions to promote or prevent EMT in clinical settings. Namely, Wnt signaling and many of the core transcription factors had notable effects on the distribution of states when perturbed. Combinatorial gene knockdowns may magnify these effects, as many feedback mechanisms exist within the EMT network. In our more robust analysis of Wnt perturbation, we found that Wnt is an important inducer of EMT and a stabilizer of a hybrid EMT.

Incorporating both literature and experimental data in the study of GRNs is a strategic approach for maximizing the relevance of the network not only to the general biological process under study, but also to the specific context in which the process is observed. The nuances of cellular self-regulation can thus be explored much further than previously possible with a given dataset, as scRNA-seq allows for researchers to explore behavioral variations across and within tissue types whereas previously a single GRN would be constructed to explain a phenomenon regardless of its context.

The advantage of combining published results with experimental data is, however, limited by the quality and quantity of available scRNA-seq data. Bulk-cell RNA-seq is insufficient in its granularity to thoroughly investigate a heterogeneous dataset and due to its novelty, scRNA-seq remains relatively challenging and expensive. Additionally, scRNA-seq is limited in its accuracy and may miss important genes entirely. Due to the small sample size, it is possible that relevant aspects of the EMT network were excluded from this analysis, although the use of three embryos at three developmental timepoints mitigates this risk. Moreover, studies have investigated the role of microRNAs in regulating EMT [4,9] but due to the nature of the dataset they were not included here. RACIPE is able to simulate regulatory relationships including microRNA, however, and could be used in combination with other experimental approaches to obtain a fuller perspective.

Beyond EMT, this approach could be employed to gain an understanding of the underlying network topology of any process of interest in a specific biological system. In cases where regulatory relationships are significantly different from one case to another, as in cancer, this method could shed light on unique aspects of the system under study and generate new hypotheses to test experimentally. Studies of cancer and especially developmental processes would also benefit from time-series data, which could help understand the changing phenotypic landscape during the course of a particular process. For example, the methodology developed here could be applied with time-series data from tumor cells to track the epigenetic shifts which promote EMT during metastasis and compare these across cancer types; RACIPE could then be further applied to simulate perturbations and identify ways to target EMT.

Regulatory interactions outside of the scope of transcriptional activation and inhibition, including those governed by competitive binding sites, posttranslational modifications, and DNA accessibility are further nuances that escape this analysis and could better illuminate the mechanics of EMT or any other process. However, using experimental methods like ChIP-seq and mass spectrometry, this methodology can be adapted to incorporate these types of interactions as well.

Here we have developed a GRN to reflect the behavior of EMT in the specific context of the embryonic mouse, identifying both interactions which regulate EMT universally and interactions which may be tissue-specific. The GRN construction protocol integrates literature-based networks and single cell transcriptomics to construct an accurate model of a particular dataset. In the case of EMT, we identified a hybrid phenotype in the scRNA-seq data as well as the simulation results and characterized the behavior of the network in response to multiple perturbations. This approach could also be used to unveil the regulatory mechanisms of a wide range of biological processes by producing in silico models which closely mirror the behavior of an experimental dataset.

## Methods

### Processing expression data and inferring transcription factor activity

We first analyzed public scRNA-seq data from 1916 cells in eight tissues of three mouse embryos at three different developmental stages ranging from E9.5-E11.5 [5]. Beginning from the log2(TPM/10 + 1) expression matrix, genes expressed in less than 1% of the cells and genes with a read count below 3% of the number of cells were removed from the dataset. SCENIC was used to infer the major transcription factors and their activity using gene expression data and regulator-target relationships from RcisTarget [19]. The algorithm infers co-expression modules using GRNBoost2 and, for each TF, identifies the direct targeted genes (*i.e.*, a regulon of the TF) with corresponding annotations in genome ranking databases. Only regulons with RcisTarget motif enrichment scores above a threshold of 3 were kept. Cells were then scored for the activity of each regulon with AUCell, yielding a regulon activity matrix [40,41]. After all 1916 cells were processed with SCENIC, we analyzed a subset of 156 skin cells independently because the subset provided more robust intermediate states.

To identify the main regulatory changes across phenotypes in the dataset, differences in regulatory link activity between clusters were evaluated using the regulon activity matrix provided by SCENIC. For each cell type cluster, the mean activity level of each regulon was calculated. The regulons with the greatest difference in mean activities between clusters were selected shown on a heatmap with the ComplexHeatmap package in R, using Spearman correlation distance and the Ward hierarchical clustering method [42–44].

### Identifying differentially expressed genes and transcription factors

For our differentially expressed genes analysis, the expression data values were normalized to a mean of 0 and genes with 0 variance were removed. We employed the Seurat package to detect DEGs between two or more given clusters, specifically using the FindAllMarkers method and the “roc” test [40,41]. In addition to identifying DEGs, several ranking scores including average log fold change and cluster classification power were generated, which were later used to rank genes and examine the activity of KEGG signaling pathways in the dataset.

### Identifying hybrid states from gene expression

We separated E and M cells by principal component analysis (PCA) across the entire filtered set of 16082 genes and all cells of a tissue type, followed by density-based clustering using the HDBClust package. These identities were consistent with the designations from [5]. To generate a list of E and M markers, DEG analysis was performed on the two clusters. The dataset was then split into subgroups of E and M cells before identifying hybrid phenotypes. For each cell type, hierarchical clustering was performed on the top 25 markers for the opposite cell type as identified by DEG analysis. Euclidean distance and the Ward.D2 clustering method were used to cut each dataset into two clusters, and the cells expressing markers of the opposite type were labeled as E-Hyb or M-Hyb according to their initial classifications.

### Network model construction

When filtering out genes with consistently low expression, nodes for which ≥80% of the cells showed expression values below the 10^th^ percentile of expression values for that gene were removed from the network. The same cutoff was applied to remove low-activity TFs from the network using the regulon activity metric in place of expression values.

For each of the interactions derived from SCENIC, a correlation score was calculated between the expression of the target and source genes, as well as the regulon activity of the source and target genes if both were TFs. Among the set of interactions suggested by SCENIC, those for which both the activity and expression correlations across skin cells were below 0.6 were removed. Because this scheme altered the network topology, genes which became inputs or outputs of the system were removed as well. After some manual adjustment to remove topologically redundant genes, meaning those which shared the same set of ≤3 interactions, the final network used in simulations contained 14 nodes and 34 edges.

### RACIPE simulations and gene perturbations

The network models were simulated with RACIPE [9,37] for stochastic analysis, where all gene expression profiles were computed from 10,000 models with randomly perturbed kinetic parameters (using one initial condition for each model). Simulated annealing was performed with an initial noise level of 13 and a noise scaling factor of 0.5 with 30 noise levels. Noise levels for each gene were scaled according to the gene expression values. State clustering was performed using spearman correlation distance and Ward.D2 clustering. Knockdown and overexpression analyses were performed by subsetting the simulation results to only include models with production rates of a given gene in the top or bottom 10% of the parameter range.

To examine the effects of a varying Wnt signal, perturbation simulations were performed by generating fresh initial conditions and setting the production rates of the gene of interest, Wnt, to five subsets of the original parameter range in increments of 20%. Stochastic simulations were then conducted to generate 10,000 models under each of these conditions.

## Acknowledgements

The study is supported by a startup fund from The Jackson Laboratory, by the National Cancer Institute of the National Institutes of Health under Award Number P30CA034196, and by the National Institute of General Medical Sciences of the National Institutes of Health under Award Number R35GM128717.

## SI Captions

**Figure S1.** Gene expression heatmaps of E/M DEGs across tissue types. (A) Skin cells, which were selected for further analysis. Note the distinct column showing co-expression on the left side of the plot. (B) Expression heatmap of intestinal cells. (C) Expression heatmap of liver cells (D) Expression heatmap of lung cells.

**Figure S2.** Dotted plots showing the number of cells at each developmental stage and belonging to each phenotype. Size and color both reflect cell count. There is a discernible upward trend in the prevalence of E and E-Hyb cells accompanied by a decrease in M cells, but the size and nature of the sample preclude a robust analysis.

**Figure S3.** Deterministic RACIPE simulation results for the final 14-node network. (A) Heatmap of the steady state gene expression profiles from deterministic RACIPE simulation results for the final 14-node network. Models are hierarchically clustered into three groups using the Ward.D2 method and spearman distance metric. The phenotypes present are comparable to the stochastic results but differ in their respective prevalence. Additionally, there is a group of models denoted by the blue-banded cluster which show low expression for all genes in the network. (B) PCA plot of the RACIPE results in part (A), color coded by cluster.

**Figure S4.** Stochastic RACIPE simulation results for the 10-node core network comprising the core interactions and the Wnt signaling pathway (A) Heatmap of the steady state gene expression profiles from stochastic RACIPE simulation results for the 10-node network comprising the core interactions and the Wnt signaling pathway. Models are hierarchically clustered into 3 groups using the Ward.D2 method and spearman distance metric. The three phenotypes present are highly similar in composition and prevalence to the results of the 14-node network. (B) PCA plot of the simulation results from part (A), color coded by cluster.

**Table S1.** Nodes present in each iteration of the network.

**Table S2.** Edges present in the final 14-node network with references if the interaction came from literature.

**Table S3.** Numbers of recorded experimental results finding the network genes in the tissues of the embryonic mouse. Results drawn from the Gene Expression Database (http://www.informatics.jax.org/expression.shtml) Ambiguous findings are recorded as such.

**Table S4.** Top 10 signaling pathways identified by fgsea GSEA (fgsea) sorted by adjusted p-value.

