## Supplementary figures and images for "Modeling a gene regulatory network of EMT hybrid states for mouse embryonic skin cells"

### Supplemental Figures

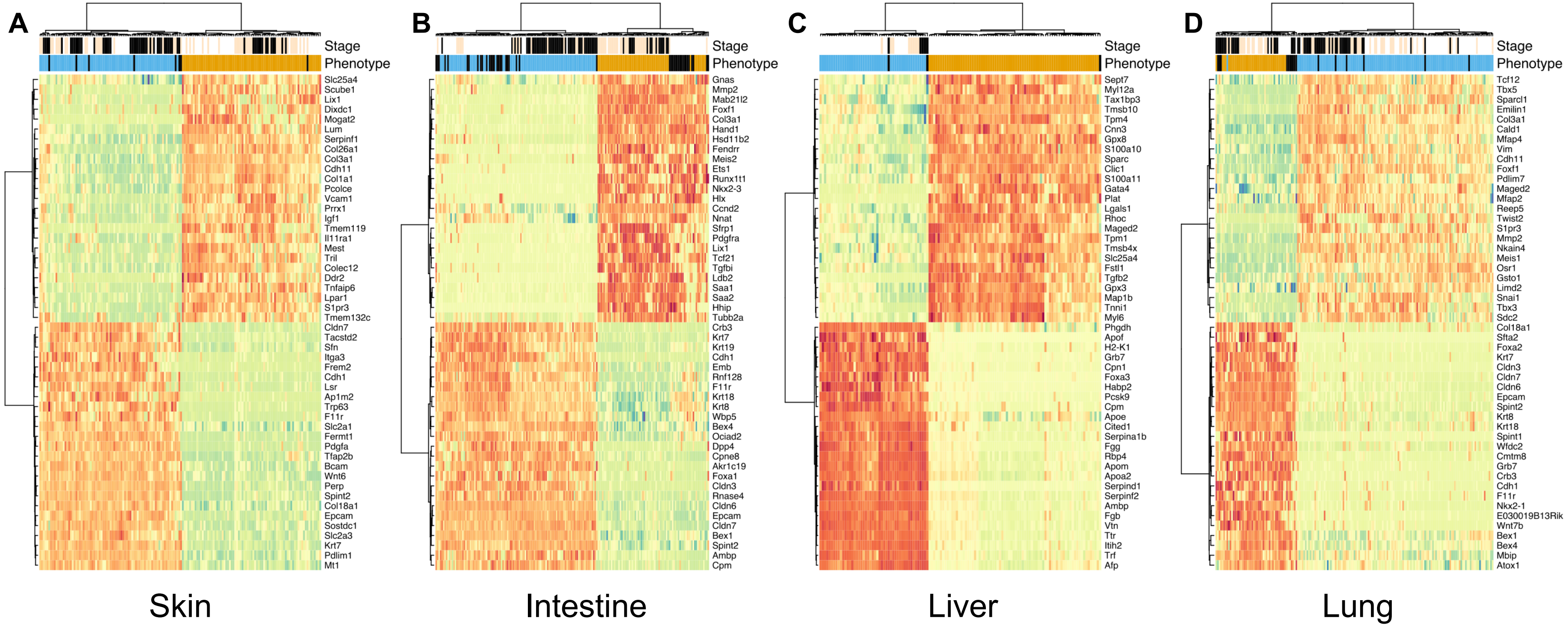

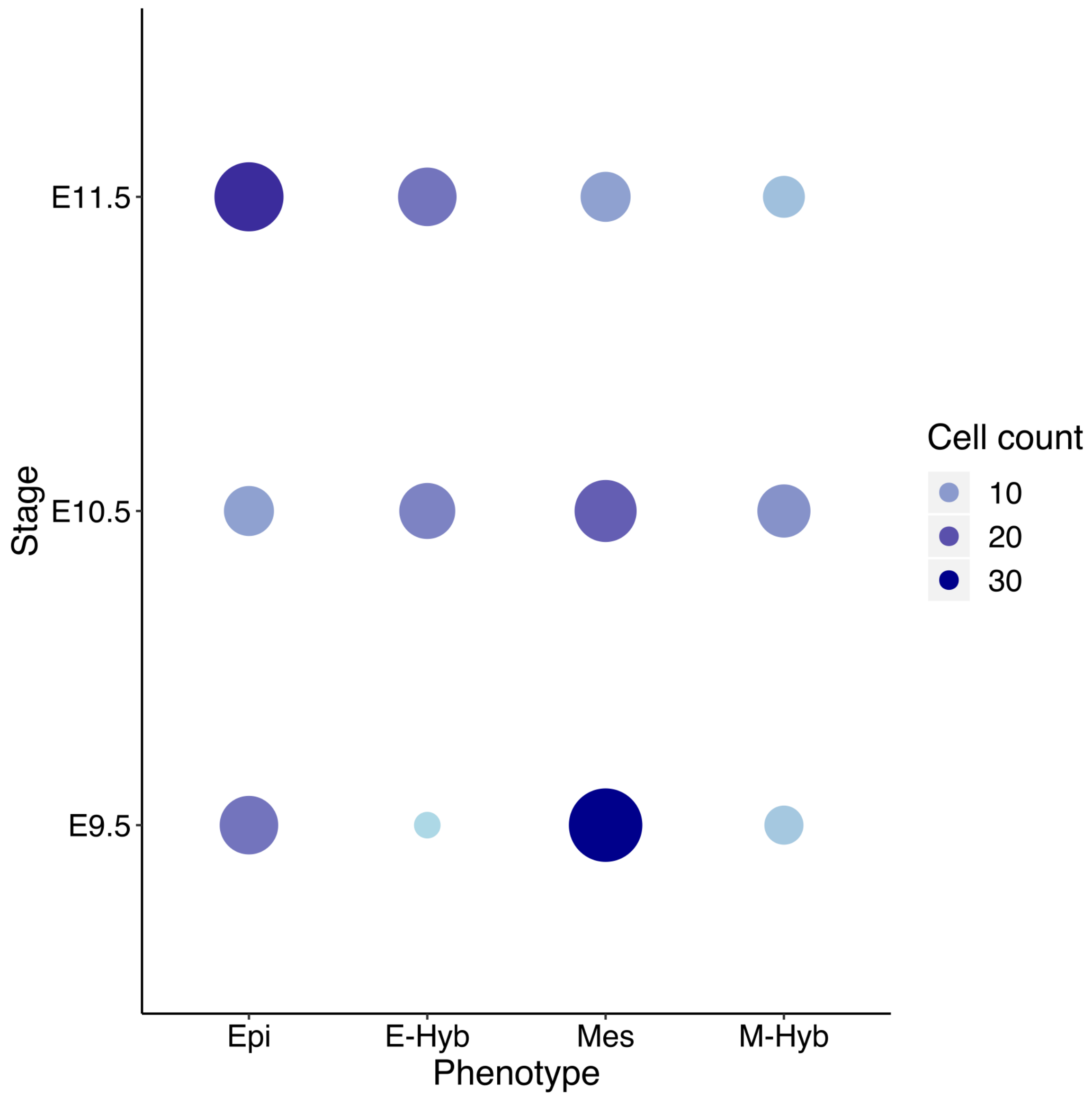

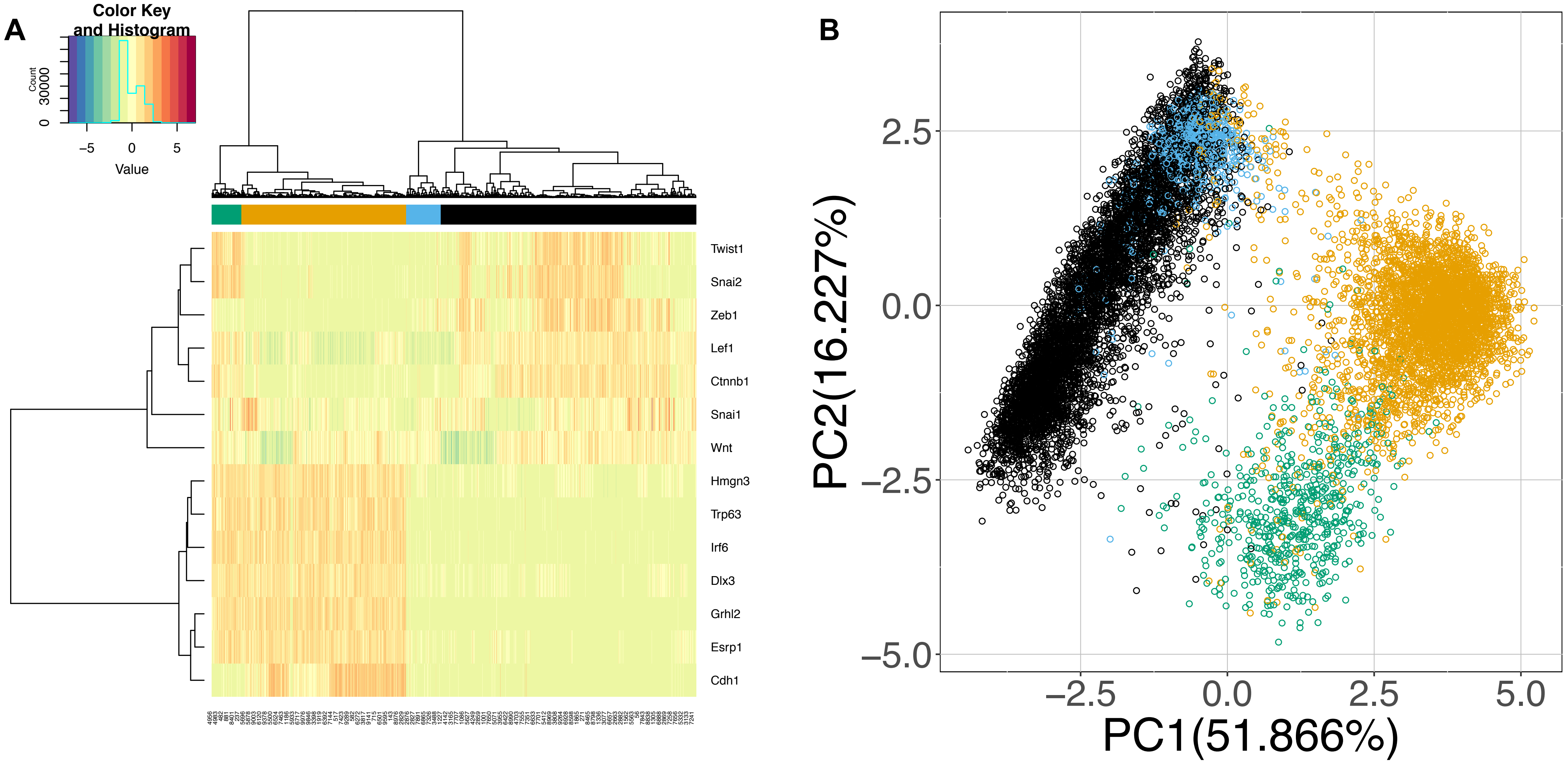

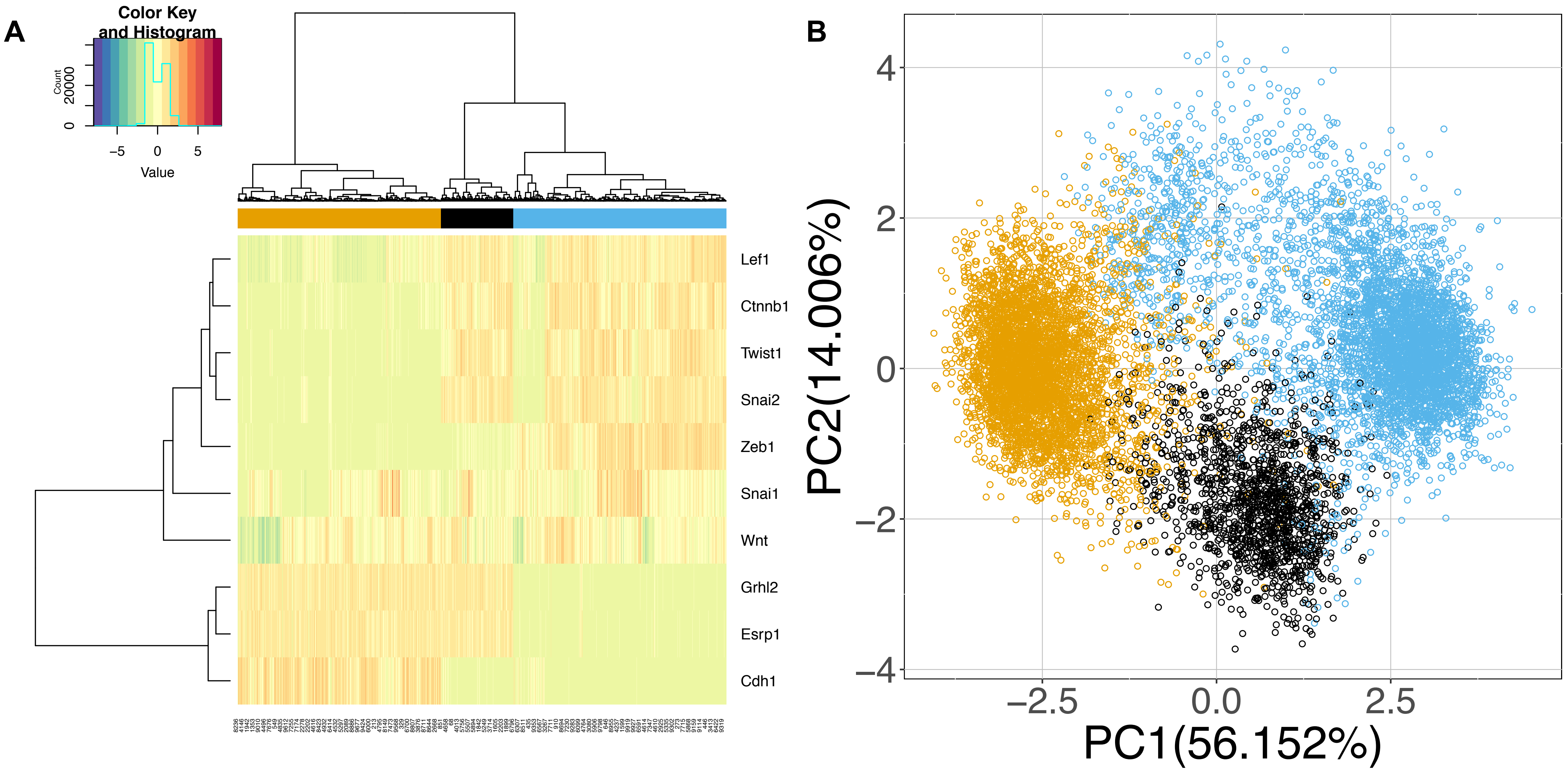
